# The role of disease state in confined migration

**DOI:** 10.1101/2024.09.16.608435

**Authors:** Annika Meid

## Abstract

Cell migration is a fundamental process in both normal and cancerous tissues, playing a crucial role in development, immune responses, and, in the case of cancer cells, metastasis-a leading cause of cancer-related mortality. Understanding the differences between healthy and cancerous cell migration is essential for uncovering potential therapeutic targets. This study aims to elucidate these differences by comparing the migratory behaviors of healthy cells (nHDF cells) and cancer cells (MDA-MB-231 cells). Our findings reveal that cancer cells significantly reduce their stiffness during migration through narrow channels, a phenomenon not observed in healthy cells. Additionally, DNA and membrane repair mechanisms are more active in healthy cells during migration compared to tumor cells. Notably, the use of MDA-MB-231 FUCCI cells demonstrates that the cell cycle profoundly influences cell migration under confined conditions. These insights provide a deeper understanding of the cellular mechanisms driving migration in both healthy and cancerous cells.

## Introduction

Cell migration is a fundamental process in the human body, essential for various physiological functions such as stem cell homing, immune defense, development, and wound healing [1]. However, in cancer, this process is hijacked, allowing cells to disseminate from the primary tumor, invade the bloodstream, and establish metastases in distant tissues and organs. Migration can occur in different forms, with cells either moving individually or as part of a collective, executing highly coordinated movements.

Single-cell migration can take on various modes. Amoeboid migration, characterized by a rounded cell shape, involves movement through blebbing, while mesenchymal migration features spindle-shaped cells that pull on the substrate to propel themselves [2]. Cancer cells are capable of switching between these migration modes depending on the availability of matrix metalloproteinases (MMPs) to degrade the extracellular matrix (ECM) and the activity of the Rho/ROCK signaling pathway [3, 4]. When ECM degradation is inhibited, cells squeeze through pores, a process constrained by the nuclear cross-sectional area, typically limited to 10% of the nuclear diameter. This critical size limit, which varies by cell type, determines whether a cell can pass through a pore without degrading the ECM.

In vivo, cells frequently migrate along pre-existing tracks within tissues, often guided by gradients of signaling molecules. For example, during wound healing, injured cells secrete TNF-α, creating a gradient that immune cells can track and follow. Coordinated migration is crucial for proper wound closure, with cells communicating through junctions and migrating as a collective, often resembling a sheet or stick-like formation [5, 6]. This mode of migration has also been observed in tumor cells, which can transition between collective and single-cell migration.

To study cell migration, various model systems have been developed that mimic the natural paths cells encounter in tissues. These include micro-patterned geometries, collagen or PEG gels, and microfluidic devices that simulate blood flow, influencing cell behavior. Microchannels, in particular, are used to replicate processes like intravasation into the bloodstream and extravasation into new tissues [4, 7, 8]. These models have become invaluable tools for understanding the complex dynamics of cell migration in both healthy and pathological contexts.

A critical step in tumorigenesis is the epithelial-to-mesenchymal transition (EMT), during which cells alter their signaling pathways and migration modes to invade surrounding tissues [9]. This transition is characterized by the upregulation of degrading enzymes, such as matrix metalloproteinases (MMPs), which facilitate infiltration into the blood or lymphatic systems. Concurrently, the disruption of communication through adherens junctions leads to the dissemination of cells from the primary tumor or cell clusters, resulting in single-cell migration [10].

The nucleus is a key limiting factor during cell migration, as it is the largest and most rigid component of the cell [11-13]. Nuclear stiffness is primarily regulated by the expression of lamin A/C. Reduced lamin A/C levels result in more deformable nuclei, which, while enhancing migration speed and efficiency, also increase the risk of nuclear damage. When cells migrate through narrow pores, the nucleus can rupture, leading to an exchange of intranuclear and cytoplasmic proteins [14, 15]. These ruptures are marked by the recruitment of proteins from the endosomal sorting complex required for transport (ESCRTIII), including charged multivesicular body proteins (CHMPs). Specifically, CHMP4B proteins are localized to the damaged nuclear envelope, playing a critical role in membrane repair [16].

During nuclear rupture, DNA double-strand breaks (DSBs) can occur, posing a significant threat to cell viability [17, 18]. In the G1 phase, DSBs are typically repaired through non-homologous end joining (NHEJ), a process in which the 53BP1 protein is recruited to the DSB site to facilitate DNA end fusion [18]. In other cell cycle phases, homologous recombination (HR) is utilized for repair, as an identical copy of the damaged DNA region is available. HR is generally error-free, with the 53BP1 protein being antagonized and prevented from binding to the damage site. If the damage is irreparable, the cell may enter senescence to prevent further cell cycle progression and potential division. This senescence is often induced in aged cells [19]. A hallmark of tumor cells is their ability to bypass these regulatory mechanisms, enabling continuous division and growth. This is often achieved through mutations in critical proteins like retinoblastoma protein or p53, which are commonly found in cancers [20-22].

Cell area and volume change during the cell cycle, leading to faster migration in the G1 phase compared to other phases. The cell cycle is intricately involved in numerous cellular processes, including adhesion formation, chromatin condensation, and DNA repair mechanisms [23-26]. It has hypothesized that tumor cells may acquire enhanced metastatic potential due to DNA damage incurred during migration. In this study, we demonstrate that highly metastatic tumor cells (MDA-MB-231) exhibit distinct behaviors compared to non-metastatic cancer cells (MCF-7) and healthy cells, particularly in their ability to reduce stiffness during migration through very narrow pores, thereby minimizing damage. The results highlight the close relationship between constriction size, cell cycle state, and the area and volume of migrating cells.

## Materials and Methods

### Fabrication of PDMS Chips

Polydimethylsiloxane (PDMS; Sylgard® 184 Silicone Elastomer Kit, DowCorning® GmbH) was prepared by mixing the base and curing agent in a 1:10 ratio, followed by casting into microfluidic chips. The chips were incubated for at least 6 hours to ensure proper curing. Subsequently, a rectangular hole was cut in the center of each chip for cell seeding. The wafer design was adapted from Rolli et al. [8]. Afterward, the chips were plasma-activated and bonded to glass slides (22 × 22 mm, Corning®). The glass slides were incubated at 80 °C for 30 minutes to reinforce the bond. The chips were then coated with 60 µg/mL collagen I (Rat tail, Gibco) and stored overnight at 4 °C. The following day, the collagen solution was removed, and phosphate-buffered saline (PBS) was added to the chips, which were stored at 4 °C until use.

### Cell culture

The breast cancer cell line MDA-MB-231 (ATCC® HTB-26™) and the HaCaT cell line (DKFZ) were maintained in DMEM GlutaMAX-I with pyruvate and 4.5 g/L D-glucose (Gibco), supplemented with 10% fetal bovine serum (FBS; Gibco) and 1% penicillin-streptomycin (PenStrep; Gibco). Primary dermal fibroblasts (ATCC® PCS-201-010™) were cultured in Medium 106 (Gibco) supplemented with the complete LSGS Kit (Gibco), resulting in final concentrations of 2% FBS, 1 µg/mL hydrocortisone, 10 ng/mL human epidermal growth factor (hEGF), 3 ng/mL basic fibroblast growth factor (bFGF), 10 µg/mL heparin, and gentamicin/amphotericin as antibiotics. The MCF-7 breast tumor cell line (ATCC® HTB-22™) was cultured in the same medium as the MDA-MB-231 cells, with the addition of 1% non-essential amino acids (NEAA; Gibco). Cell counts were performed using a Beckman Coulter Counter (Beckman AG 4438).

### Isolation of the plasmid nls-eGFP

The nls-eGFP plasmid (Addgene plasmid 67652), designed by Rob Parton, was delivered in E. coli DH5α cells. The bacteria were grown overnight on LB agar plates containing 100 µg/mL ampicillin using the three-streak method. A single colony was then inoculated in LB medium with 100 µg/mL ampicillin and cultured overnight at 200 rpm and 37 °C. Plasmid DNA was isolated using the Qiagen Plasmid Midi Kit, and the final concentration was measured using a Tecan spectrophotometer.

### Transfection of the cell lines

MDA-MB-231, MCF-7, and HaCaT cells were transfected with the nls-eGFP plasmid using Lipofectamine® 2000 (Invitrogen). For each transfection, 500 ng of plasmid DNA was combined with 1.4 µL Lipofectamine. nHDF cells were transfected using the Amaxa nHDF Nucleofector™ Kit (Lonza) and the U-020 program for human neonatal HDF with the Nucleofector™ 2b Device (Lonza).

### Seeding of the cell into PDMS chips

PDMS chips were sterilized under UV light for 1 hour before cell seeding. Approximately 1 × 10⁵ to 2 × 10⁵ cells were seeded into the center of each chip. After initial attachment was confirmed, an additional 2–5 mL of medium was added to fully cover the chip. Cells were maintained at 37 °C and 5% CO₂ until the migration process through the channels commenced. Cells were either fixed or imaged over several days. MDA-MB-231 FUCCI cells (gifted by Prof. Bojana Gligorijevic) were seeded at a density of 2 × 10⁵ cells per chip. After attachment, the chips were covered with medium and imaged over the following 2–3 days.

### Antibodies and staining of fixed cells

For cell fixation during migration through the channels, cells were treated with 4% paraformaldehyde (PFA) for 10 minutes. Permeabilization was performed using 0.1% Triton X-100 in PBS (Roth) for 15 minutes. Cells were stained for CHMP4B (1:100, rabbit, Abcam, ab105767) and Lamin A/C (1:100, mouse, Santa Cruz Biotechnology, Sc-7292) or with Lamin A/C and 53BP1 (1:100, rabbit, Invitrogen, PA1-16565) in PBS-T with 1% BSA (PAA). The primary antibodies were incubated for 1 hour at 37 °C, followed by washing with PBS. Secondary antibodies were then applied, including Hoechst 33342 (1:500, Invitrogen H1399I), AlexaFluor™ 568 Phalloidin (1:500, Invitrogen, REF A12380), AlexaFluor™ 488 Chicken anti-rabbit IgG (1:500, Invitrogen, REF A21441), and AlexaFluor™ 647 goat anti-mouse IgG (1:500, Invitrogen, REF A21235). The chips were incubated for 30 minutes at 37 °C in a 5% CO₂ atmosphere. After washing with distilled water, the chips were glued to Petri dishes with Picodent twinsil® and stored at 4 °C until imaging.

### Microscopy and data analysis

Cell volume measurements were performed by imaging z-stacks using an LSM980 confocal microscope (Zeiss). Transfected cells, MDA-MB-231 FUCCI cells, and fixed cells were imaged using either an Axiovert 200M or Axioobserver Z1 microscope. Intensity measurements were analyzed using CellProfiler, while all other analyses were conducted using Fiji. Statistical comparisons were made using Welch’s t-test, with results presented as 95% confidence level (CL) error bars. All graphs were generated using Prism and Adobe Illustrator. All experiments were performed at least three times.

### Results and Discussion

To assess the nuclear properties of cells during migration, fixed cells were stained and analyzed for various parameters, including aspect ratio (AR) and nuclear area. The AR was calculated for cells before, during, and after migration through microchannels of different widths (Figure 1).

**Figure 1:**
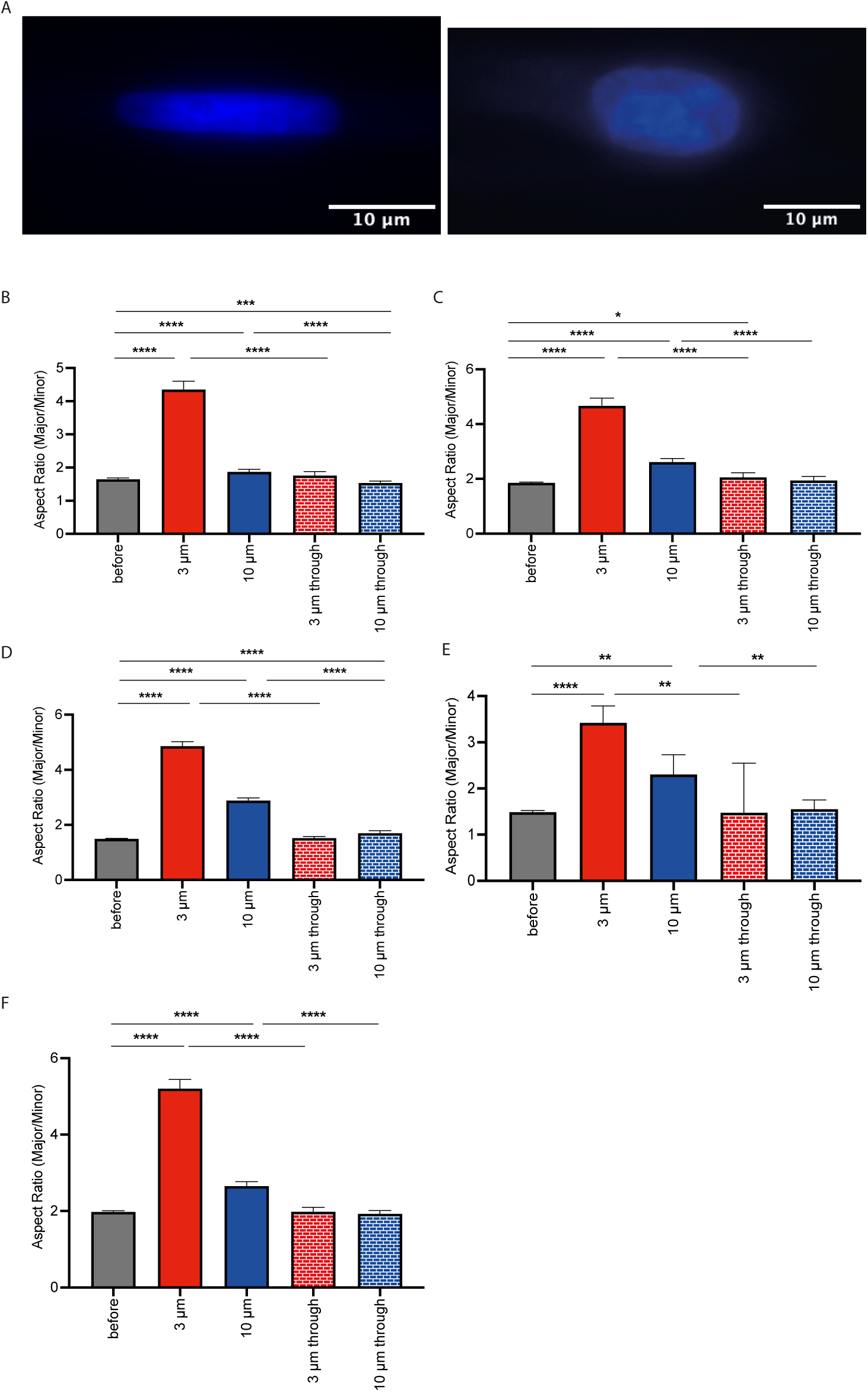
Aspect ratio analysis of various cell lines before, during, and after migration through narrow channels. (1A) Left: Hoechst-stained nucleus of an MDA-MB-231 cell within a 3 µm channel. Right: Hoechst-stained nucleus of an MDA-MB-231 cell within a 10 µm channel. (1B) Aspect ratio of MDA-MB-231 cells before migration, during migration through 3 µm and 10 µm channels, and after migration. *n=9*. (1C) Aspect ratio of nHDF cells under the same conditions. (1D) Aspect ratio of HaCaT cells under the same conditions. (1E) Aspect ratio of MCF-7 cells under the same conditions. (1F) Aspect ratio of HFF cells under the same conditions. The maximum number of cells was analyzed for each cell line and condition. Data were analyzed using Prism software, and statistical comparisons were performed using Welch’s t-test. Error bars represent 95% confidence levels (CL).

Figure 1A illustrates the distinct shapes and ARs of MDA-MB-231 cell nuclei within 3 µm and 10 µm channels. MDA-MB-231 cells, a highly aggressive tumor cell line, exhibited the highest AR within 3 µm channels and the lowest AR within 10 µm channels, with AR values slightly elevated compared to non-migrated cells (Figure 1B). This suggests that channel width significantly influences nuclear elongation during migration, a trend observed across all cell lines tested. For instance, fibroblasts (nHDF) demonstrated higher ARs in 10 µm channels (Figure 1C). Similarly, HaCaT and MCF-7 cells displayed a significant increase in AR within 10 µm channels, with a further increase in narrower channels (Figures 1D and 1E). Importantly, after migrating through the channels and re-entering non-confining conditions, AR values returned to baseline levels comparable to those of non-migrated cells, as observed in HFF cells as well (Figure 1F).

To determine whether nuclear size limits entry and passage through the channels, nuclear area measurements were performed. Consistent with findings by Wolf et al., cells are required to degrade the extracellular matrix if more than 10% of the nuclear cross-section enters a channel—a process impeded in stiff PDMS gels. In MDA-MB-231 cells, nuclear area decreased significantly within narrow 3 µm channels (Figure 2A), while no difference in nuclear area was observed in wider channels when compared to control cells. Notably, cells that passed through the 3 µm channels maintained a reduced nuclear area even post-passage. In contrast, healthy nHDF cells exhibited a substantial reduction in nuclear area across all channel widths, with a complete recovery of nuclear area after channel passage (Figure 2B). This reduction may be attributed to cell shrinkage caused by water efflux. Similar reductions in nuclear area were observed in HaCaT, MCF-7, and HFF cells across all channel widths (Figures 2C, 2D, and 2E). These results suggest that only MDA-MB-231 cells can pass through narrower channels without significant nuclear shrinkage, whereas healthy and non-metastatic cells must reduce their nuclear area to navigate these constraints.

**Figure 2:**
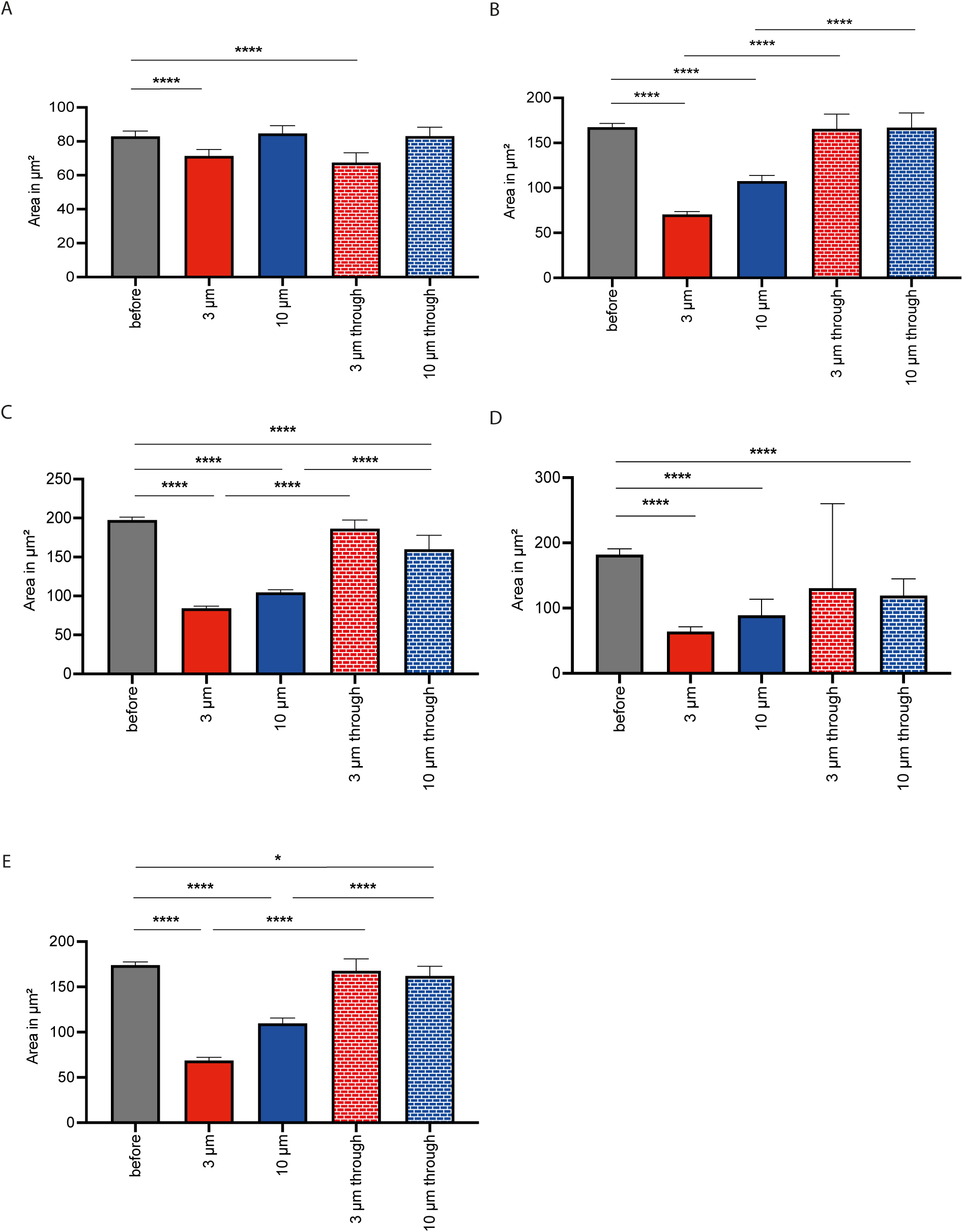
Nuclear area measurements of different cell lines before, during, and after migration through microchannels. (2A) Nuclear area of MDA-MB-231 cells before migration, during migration through 3 µm and 10 µm channels, and after migration. *n=9*. (2B) Nuclear area of nHDF cells under the same conditions. *n=8*. (2C) Nuclear area of HaCaT cells under the same conditions. *n=6*. (2D) Nuclear area of MCF-7 cells under the same conditions. *n=7*. (2E) Nuclear area of HFF cells under the same conditions. *n=13*. The maximum number of cells was analyzed for each condition. Data were analyzed using Prism9, and significance was assessed using Welch’s t-test. Error bars represent 95% CL.

The question arose as to whether cells compensate for the narrow width of 3 µm channels by optimizing their use of channel height, effectively filling the channel volume. Our results confirm that nuclear changes are more pronounced in narrower channels, a phenomenon consistent across all tested cell lines and corroborated by previous studies.

Further analysis using confocal microscopy was conducted to explore potential differences in nuclear volume. MDA-MB-231 cells exhibited a reduced nuclear volume within both 3 µm and 10 µm channels, with the volume remaining diminished even after migration (Figure 3A). In contrast, nHDF cells were able to restore their nuclear volume to control values after migration (Figure 3B), although a reduction in volume was observed during migration, particularly within 3 µm channels (Figure 3C). HaCaT cells also displayed a lower nuclear volume during migration through 3 µm and 10 µm channels. While there was an increase in volume post-migration, it did not fully return to control levels (Figure 3D). Similarly, MCF-7 cells showed a reduction in nuclear volume within both channel widths. The nuclear volume remained reduced in cells within 3 µm channels, but was restored to control levels in cells that passed through 10 µm channels. Video examples of nHDF cells before and during migration through microchannels are provided in Videos S1 and S2.

**Figure 3:**
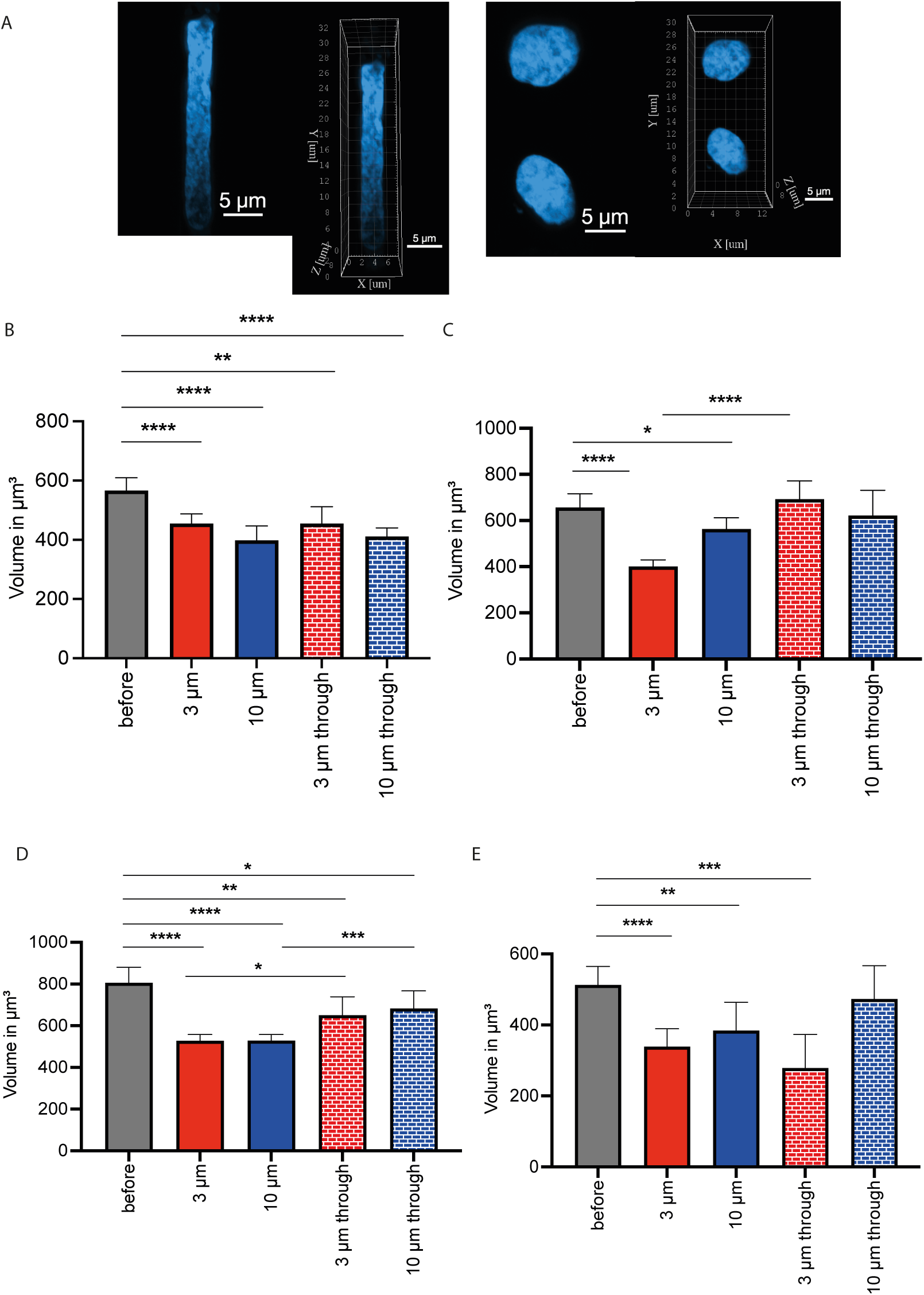
Volume analysis of cells before, during, and after migration through different channel widths. (3A) LSM 980 images of MDA-MB-231 cells within 3 µm (left) and 10 µm (right) channels. (3B) Nuclear volume of MDA-MB-231 cells in 3 µm and 10 µm channels. (3C) Nuclear volume of nHDF cells in 3 µm and 10 µm channels and after migration. (3D) Nuclear volume of HaCaT cells under the same conditions. (3E) Nuclear volume of MCF-7 cells under the same conditions. The maximum number of cells was analyzed for each condition. Data analysis was performed using Prism9, with Welch’s t-test employed for statistical comparisons. Error bars represent 95% CL.

The findings indicate that healthy cells exhibit a reduced area and volume when migrating through constricted channels, suggesting that only smaller cells can successfully pass through. For MDA-MB-231 cells, this behavior is only observed in the most narrow channels. This volume reduction during migration could be a strategy to minimize cellular damage, as seen in glioblastoma cells that shrink due to water loss during migration [27, 28]. Similar behaviors have been reported in other cancer types [29], though this response is cell type-specific [30]. Post-migration, cells partially regain their volume but remain smaller than control cells, implying potential transcriptional changes that persist at the time of fixation. To fully understand these dynamics, further studies using live-cell imaging over extended periods are necessary to determine whether cell volume can be fully restored. Additionally, staining for water pumps to detect water efflux in this model could yield valuable insights.

Previous studies have reported increased cellular damage when migrating through constricted environments [14, 31]. This prompted an investigation into whether cells could adapt to nuclear constriction during migration through microchannels where no external pressure is applied, allowing cells time to pass through. DNA and nuclear membrane damage were assessed via staining. Microscopy images in Figure 4A show increased intensity within the 3 µm channel (upper image) compared to the 10 µm channel (lower image). In MDA-MB-231 cells (Figure 4B), the intensity of the DNA repair marker 53BP1 is reduced in the smaller 3 µm channels, indicating less active DNA repair compared to control cells. However, after migrating through the shorter channels, 53BP1 intensity increases, surpassing levels observed in control cells. In contrast, nHDF cells (Figure 4C) show increased 53BP1 intensity during migration through all channel widths. Post-migration, these cells exhibit similar 53BP1 activity to control cells that did not undergo migration. HaCaT cells (Figure 4D) also demonstrate higher 53BP1 activity during migration through all channel widths, with elevated intensity persisting even after migration, unlike control cells. These results highlight the distinct DNA repair responses of different cell lines during migration through confined spaces.

**Figure 4:**
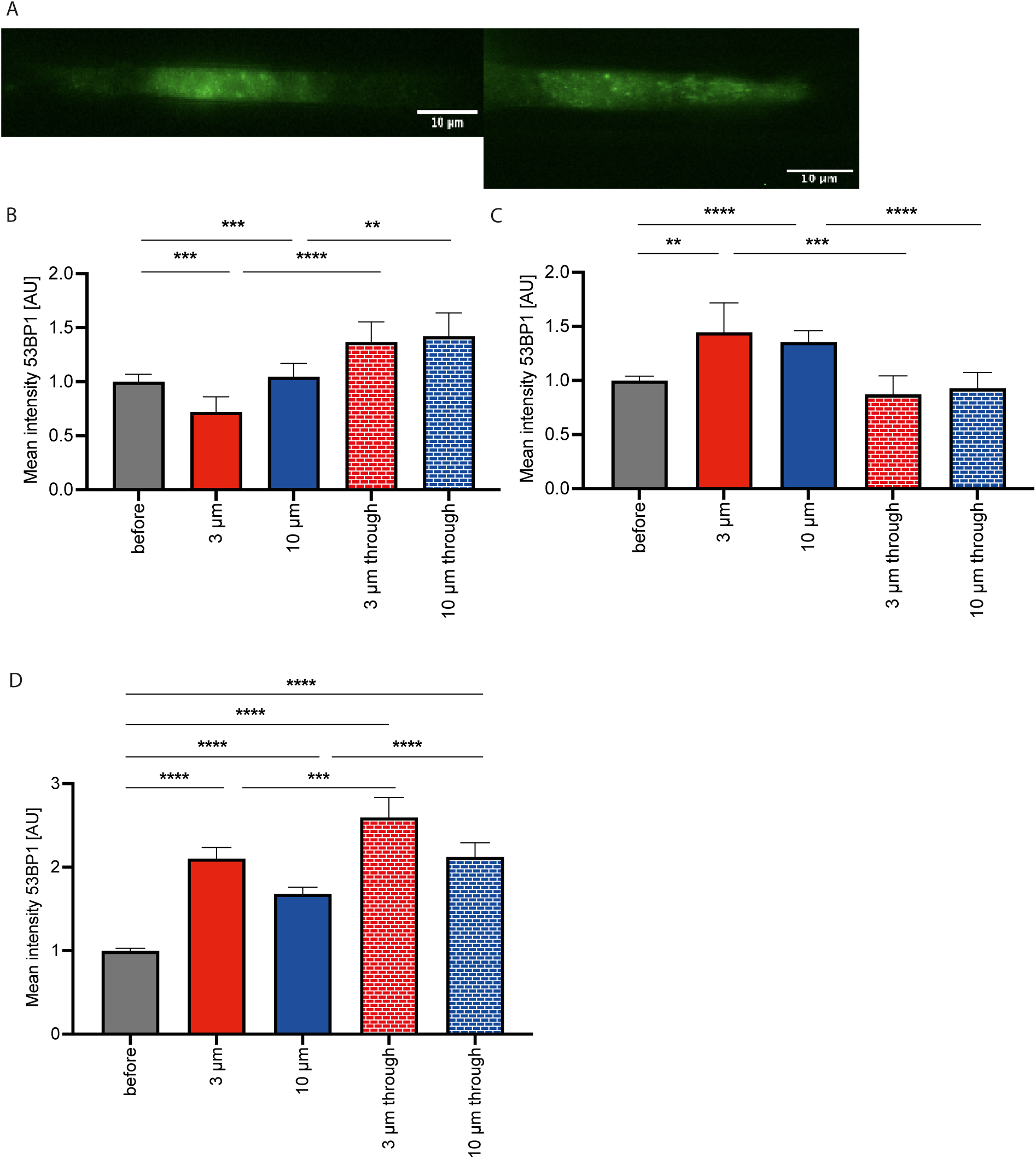
53BP1 intensity, indicative of DNA damage, in various cell lines. (4A) MDA-MB-231 cell within a 3 µm channel (top) and within a 10 µm channel (bottom). (4B) Normalized 53BP1 intensity in MDA-MB-231 cells during and after migration through microchannels. (4C) Normalized 53BP1 intensity in nHDF cells under the same conditions. (4D) Normalized 53BP1 intensity in HaCaT cells under the same conditions. The maximum number of cells was analyzed for each condition using Prism9, with Welch’s t-test used for significance testing. Error bars represent 95% CL.

Figure 5A illustrates a MDA-MB-231 cell stained for the nuclear membrane repair protein CHMP4B within 3 µm (left) and 10 µm (right) channels. MDA-MB-231 cells (Figure 5B) exhibit lower CHMP4B expression inside the channels compared to control cells, with expression decreasing as channel width narrows. After migration, CHMP4B intensity returns to levels comparable to non-migrating cells. For nHDF cells, CHMP4B intensity increases across all channel widths relative to control cells (Figure 5C). Following migration, cells that traversed the 10 µm channels exhibit CHMP4B levels similar to control cells, while those passing through 3 µm channels maintain elevated intensity. HaCaT cells (Figure 5D) show reduced CHMP4B intensity within 3 µm channels but increased expression in 10 µm channels. After migration, a higher CHMP4B intensity is observed in all HaCaT cells, regardless of the channel width they traversed.

**Figure 5:**
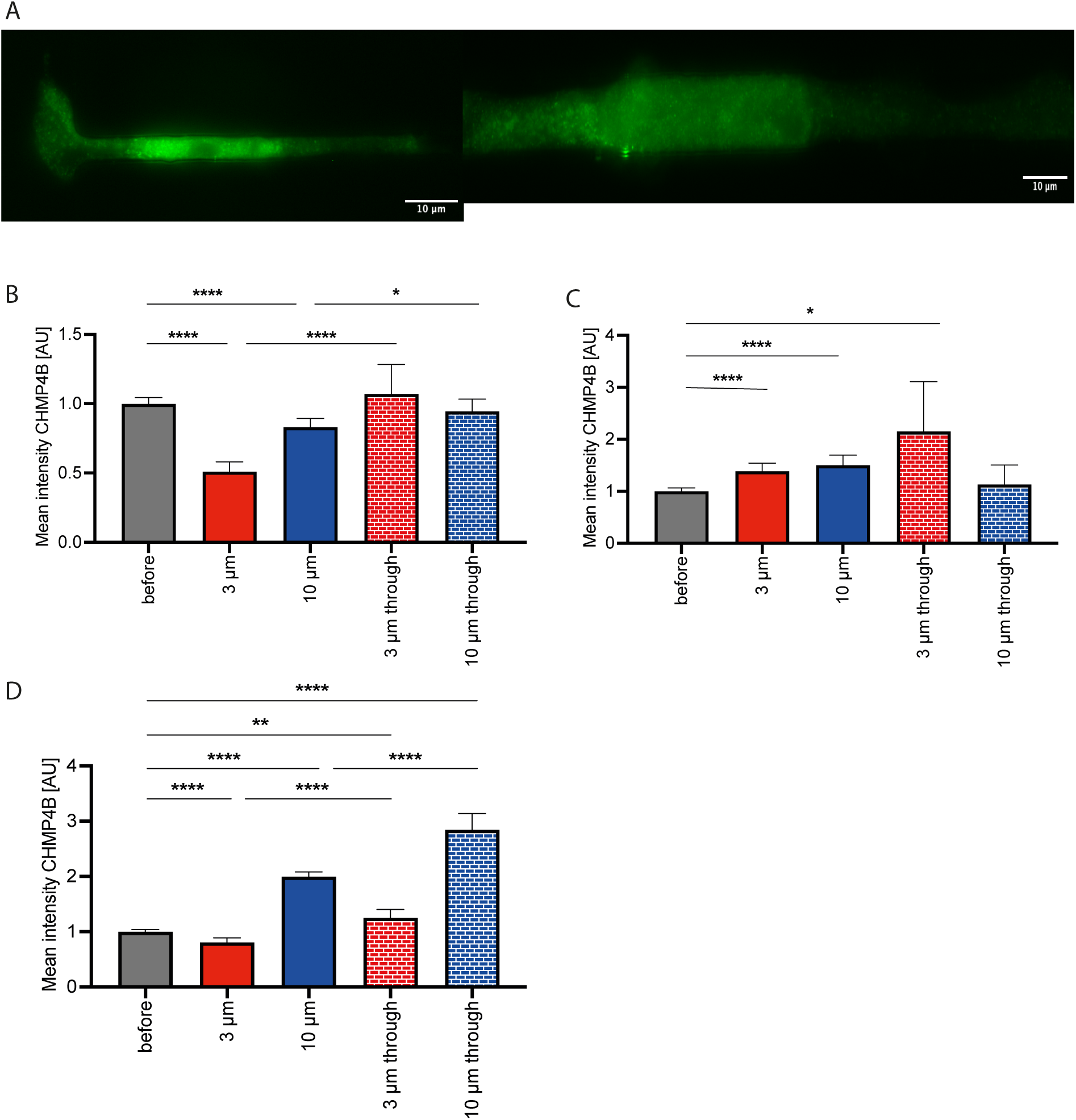
Normalized mean intensity of CHMP4B in different cell lines. (5A) MDA-MB-231 cell within a 3 µm channel (left) and a 10 µm channel (right). (5B) 53BP1 intensity in MDA-MB-231 cells during and after migration through microchannels (*n=4*). (5C) 53BP1 intensity in nHDF cells under the same conditions (*n=4*). (5D) 53BP1 intensity in HaCaT cells under the same conditions (*n=4*). The maximum number of cells was analyzed using Prism9, with Welch’s t-test applied to assess significance. Error bars represent 95% CL.

These findings suggest that tumor cells may migrate through confined spaces without adequately addressing DNA damage, potentially leading to further genetic alterations and increased tumor aggression. This observation aligns with previous studies that reported higher levels of DNA damage in tumor cells under similar conditions [14]. In this study, cells were compelled to pass through channels using microfluidic devices, highlighting the need for further research in this area to uncover novel insights. A key aspect of DNA repair is the protein 53BP1, which is critical for repair mechanisms but is often mutated in cancer cells [32-36]. Notably, 53BP1 is active only during the G1 phase of the cell cycle, while BRCA1, another protein involved in homologous recombination (HR) and active during the G2 phase, also plays a crucial role. Investigating both proteins using live-cell imaging could provide a more comprehensive understanding of DNA repair processes and their impact on cell migration under confined conditions [17, 37-39]. Membrane repair mechanisms are also impaired in cancer cells, leading to nuclear membrane rupture, which can cause DNA damage and facilitate the exchange of nuclear and cytoplasmic proteins [14, 31]. The nuclear lamina, which helps maintain nuclear integrity, prevents such ruptures and associated DNA damage, thereby preventing cell cycle arrest [40].

Cell stiffness is a critical factor in determining deformability, with reduced stiffness potentially aiding cells in migrating through narrow pores. The MDA-MB-231 tumor cell line exhibited a significant reduction in stiffness during migration through confined channels, particularly in the narrowest (3 µm) channels (Figure 6B). This reduction in stiffness, measured by lamin A/C staining, persisted even after the cells exited the channels, compared to control cells. Figure 6A illustrates a MDA-MB-231 cell inside a 3 µm channel (left) and a 10 µm channel (right), clearly showing lower lamin A/C intensity in the narrower channel. Interestingly, nHDF cells did not exhibit a reduction in stiffness; instead, they either maintained their stiffness at control levels or increased it in wider channels, especially in 10 µm channels (Figure 6C). Even after passing through the channels, the stiffness of nHDF cells remained consistent with control levels. HaCaT cells also showed a decrease in lamin A/C intensity during migration through channels of all widths, with the reduction persisting slightly after passing through 3 µm channels, while cells that passed through 10 µm channels exhibited lamin A/C levels similar to control cells.

**Figure 6:**
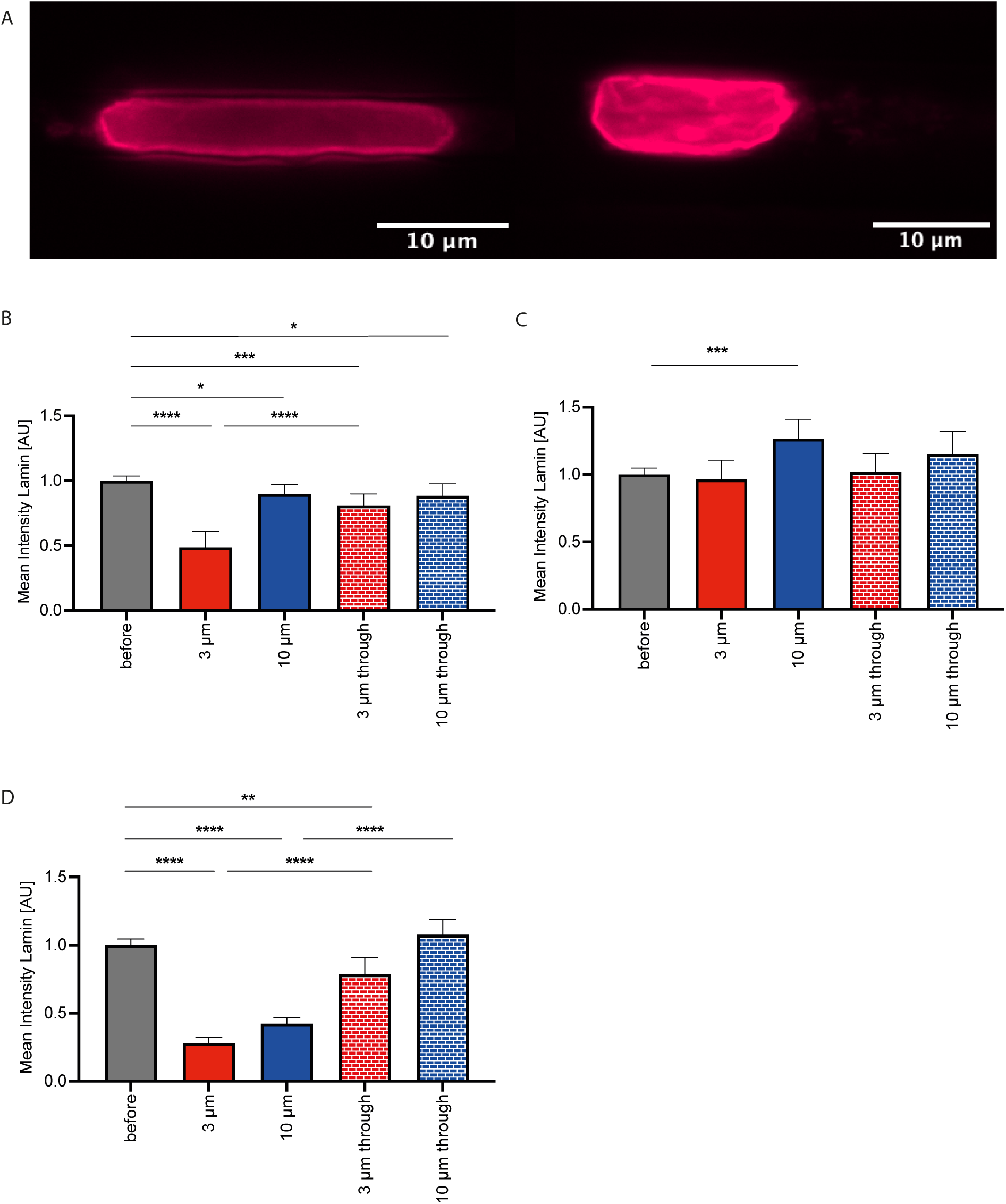
Normalized mean intensity of lamin A/C in different cell lines. (6A) MDA-MB-231 cell within a 3 µm channel (left) and a 10 µm channel (right). (6B) Normalized mean intensity of lamin A/C in MDA-MB-231 cells during and after migration through microchannels. (6C) Normalized mean intensity of lamin A/C in nHDF cells under the same conditions. (6D) Normalized mean intensity of lamin A/C in HaCaT cells under the same conditions. The maximum number of cells was analyzed using Prism9, with Welch’s t-test employed for significance testing. Error bars represent 95% CL.

These observations suggest that HaCaT and MDA-MB-231 cells reduce their stiffness to navigate through confined channels, while healthy nHDF cells tend to increase their stiffness. Lamin A/C is a primary contributor to nuclear stiffness, and its reduction leads to decreased stiffness and increased nuclear deformability. However, this reduction in stiffness is also associated with increased DNA damage [13, 14, 31] and compromised nuclear envelope (NE) integrity [41]. In MDA-MB-231 cells, the combination of reduced stiffness and diminished DNA and membrane repair supports the hypothesis that tumor cells are more susceptible to damage during confined migration, contributing to their enhanced aggressiveness and metastatic potential. Chromatin also plays a role in nuclear stiffness, migration processes, and the maintenance of DNA integrity [42-44]. Chromatin structure may serve as a marker for identifying aggressive cancers [45], and further studies are needed to explore its influence on nuclear stiffness and to uncover differences during confined migration.

Figure 7A shows two MDA-MB-231 cells migrating through a 3 µm channel (left) and a 10 µm channel (right). The cell in the 3 µm channel exhibits an elongated shape compared to the cell in the 10 µm channel. If the nuclear envelope ruptures, the signal of nls-eGFP, which is normally confined to the nucleus, can diffuse into the cytoplasm, thereby increasing cytoplasmic nls-eGFP intensity. This results in a reduction of the nls-eGFP nucleus/cytoplasm ratio, indicating higher DNA damage. This phenomenon was observed in both cell lines, with MDA-MB-231 cells showing a reduced ratio when migrating through 3 µm channels (Figure 7B). In contrast, no reduction was observed in 10 µm channels. In nHDF cells, the intensity of nls-eGFP was lower in 3 µm channels (Figure 7C). These findings indicate that the nucleus is more prone to rupture during migration through narrow 3 µm channels, leading to potential DNA damage. In larger channels, no significant nuclear breakage was observed. For the other two cell lines, HaCaT and MCF-7, only a few transfected cells successfully migrated through the channels. HaCaT cells showed no reduction in nls-eGFP intensity in 10 µm channels, and no transfected cells entered narrower channels. Similarly, MCF-7 cells did not exhibit a reduction in 10 µm channels, and no cells passed through the narrower channels. Achieving stable transfection and selecting for transfected cells could address the issue of low transfection efficiency, providing a larger population of nls-eGFP-expressing cells for future monitoring and analysis.

**Figure 7:**
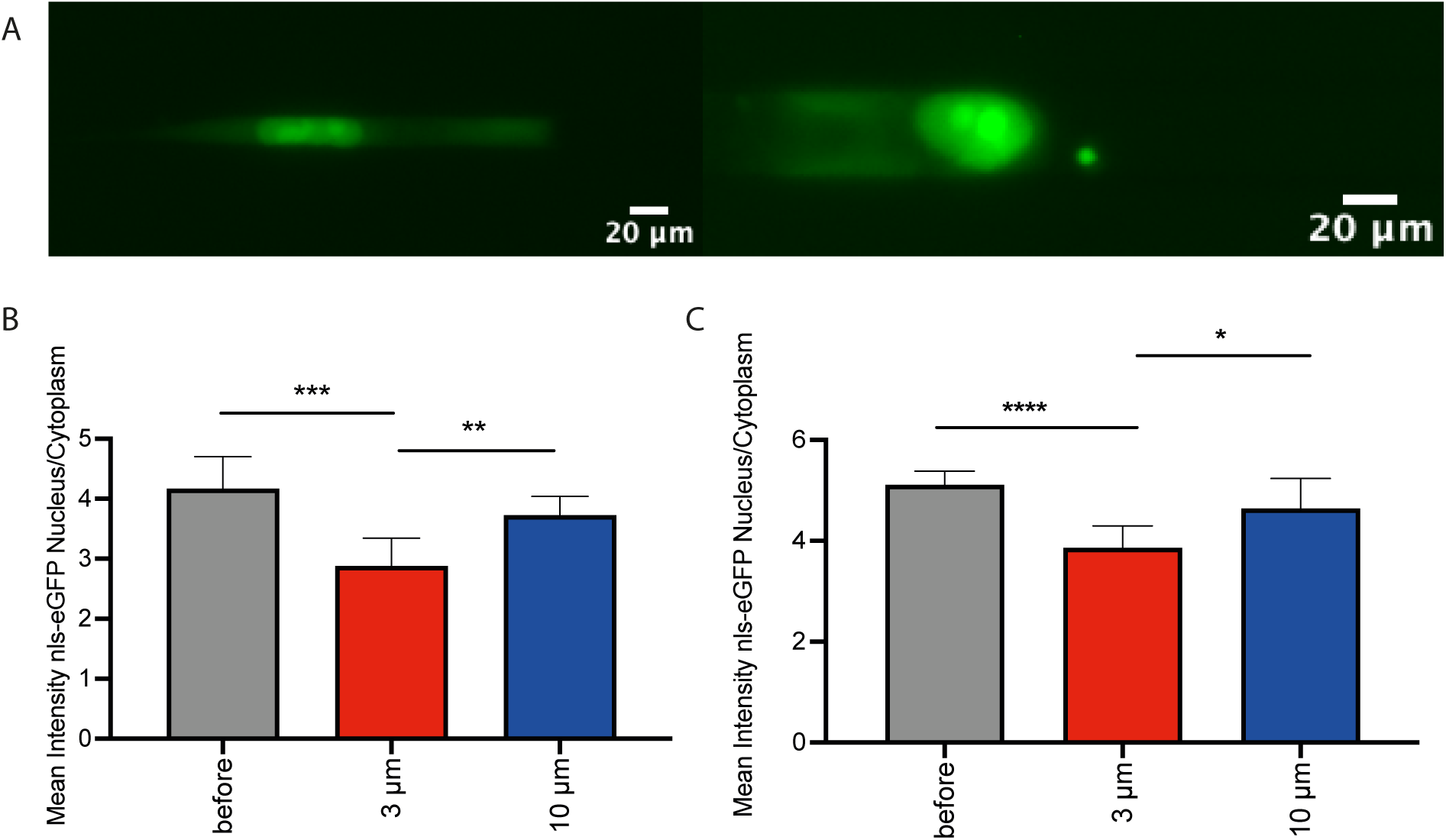
nls-eGFP intensity analysis in cells migrating through confined spaces. (7A) Left: MDA-MB-231 cell within a 3 µm channel. Right: MDA-MB-231 cell within a 10 µm channel. (7B) Mean intensity of nls-eGFP in the nucleus/cytoplasm ratio in MDA-MB-231 cells during migration through microchannels. (7C) Mean intensity of nls-eGFP in the nucleus/cytoplasm ratio in nHDF cells under the same conditions.

Live-cell experiments utilizing nls-eGFP revealed increased nuclear damage in cells migrating through 3 µm channels, as evidenced by higher nls-eGFP intensity in the cytoplasm, compared to those in 10 µm channels. This suggests that cells experience greater nuclear deformability in narrower channels, potentially leading to increased damage. However, in healthy cells, this damage may be mitigated by heightened activity of DNA and membrane repair proteins, allowing them to survive despite the stress. This observation aligns with live-cell imaging data, where healthy cells frequently blocked channels and subsequently died. The lack of such repair mechanisms in MDA-MB-231 cells within confined channels supports the hypothesis of greater nuclear damage during migration. Prior research has indicated that tumor cells exhibit a distinct mode of migration in confined spaces due to altered adhesion dynamics [4].

To investigate the impact of the cell cycle on migratory behavior, MDA-MB-231 FUCCI cells, which exhibit color changes corresponding to different cell cycle phases, were employed. As shown in Figure 8A, there was no significant time difference between the red (G1 phase) and green (late S/G2/M phase) phases. However, the early S phase (yellow) was notably shorter, with the colorless phase between M and G1 being the briefest. When migrating through 3 µm constrictions, the majority of cells were in the red G1 phase (Figure 8B), with fewer cells entering the green (late S/G2/M) and yellow (early S) phases. Transitions from yellow to green or red to green were also observed. Conversely, in 10 µm channels, a higher proportion of cells entered the green phase (late S/G2/M), with 45.16% of cells migrating through the channels in the red phase and an increase in green phase entry from 13.16% in 3 µm constrictions to 38.71% in 10 µm channels (Figure 8C). A greater percentage of cells also migrated through the channels in the early S phase, with observable transitions between G1 and late S/G2/M phases. Supplemental videos 3 and 4 provide visualizations of MDA-MB-231 FUCCI cells migrating through 3 µm and 10 µm channels.

**Figure 8:**
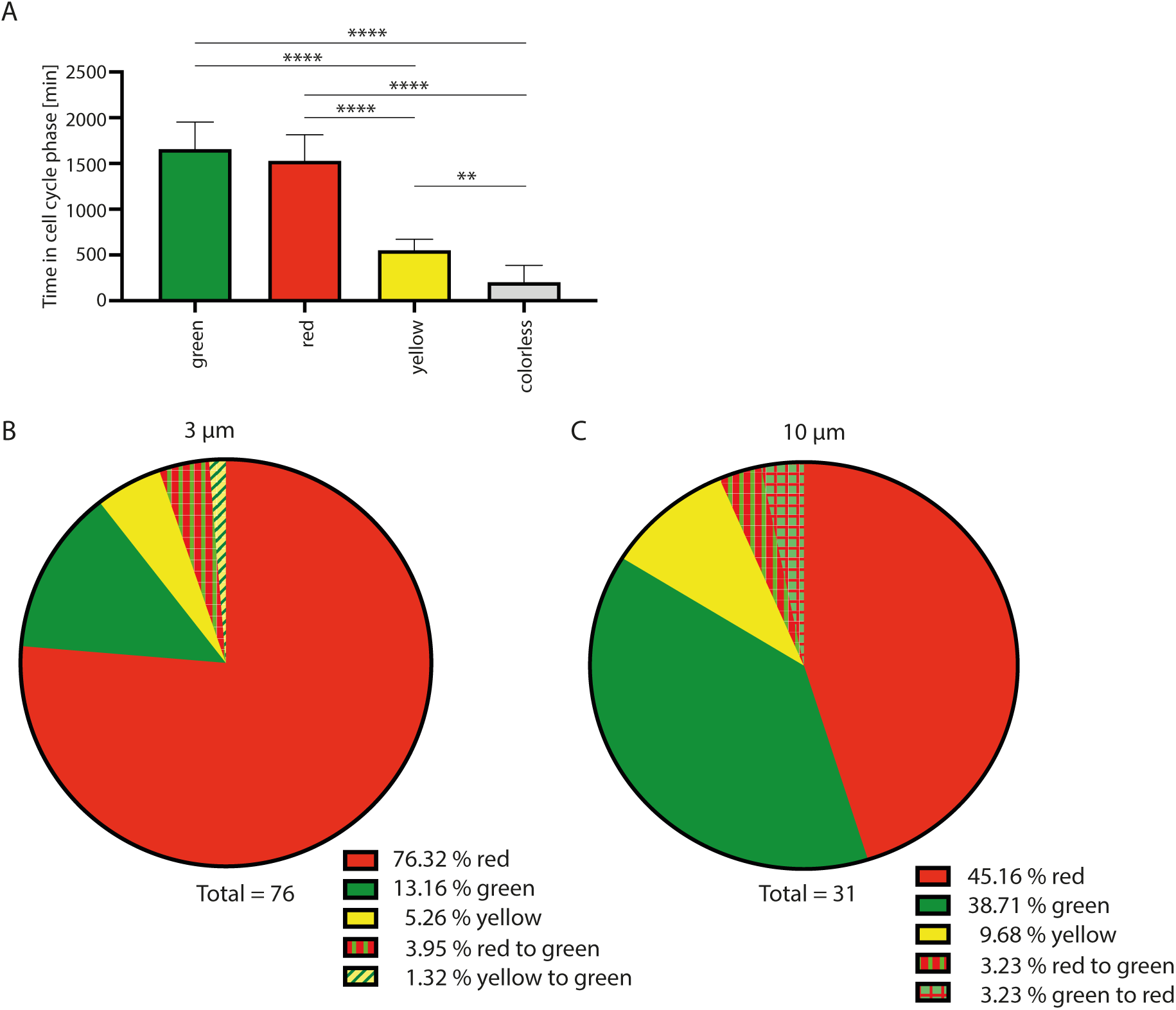
Analysis of MDA-MB-231 FUCCI cells under confinement. (8A) Duration of each cell cycle phase in MDA-MB-231 FUCCI cells. (8B) MDA-MB-231 FUCCI cells within 3 µm constrictions. (8C) MDA-MB-231 FUCCI cells within 10 µm constrictions.

These findings indicate that the cell cycle state significantly influences cell migration in confined environments, with narrower channels favoring entry into the G1 phase, where cells are more flexible and deformable. This underscores the importance of considering the cell cycle in studies of confined migration. The development of a FUCCI-labeled nHDF cell line to investigate the cell cycle’s impact on migration in healthy cells is warranted, as it is likely that these cells will exhibit similar behavior, with an even higher percentage passing through the narrowest channels during G1. The involvement of the cell cycle in processes such as cell adhesion, DNA repair, and changes in cell size and volume suggests that migration is more complex than previously understood [23-26-46-47]. Future experiments should therefore incorporate cell cycle analysis. It has been shown that cancer cells predominantly remain in the G0 and G1 phases during invasion, contributing to chemoresistance due to their non-dividing state [48].

## Acknowledgments

Prof. Bojana Gligorijevic is acknowledged for kindly providing the MDA-MB-231 FUCCI cells.

## Declaration

The authors declare no conflict of interests.

## Author contribution

AM performed all experiments and wrote the manuscript.

## Supplemental Material

**Supplemental video 1:**
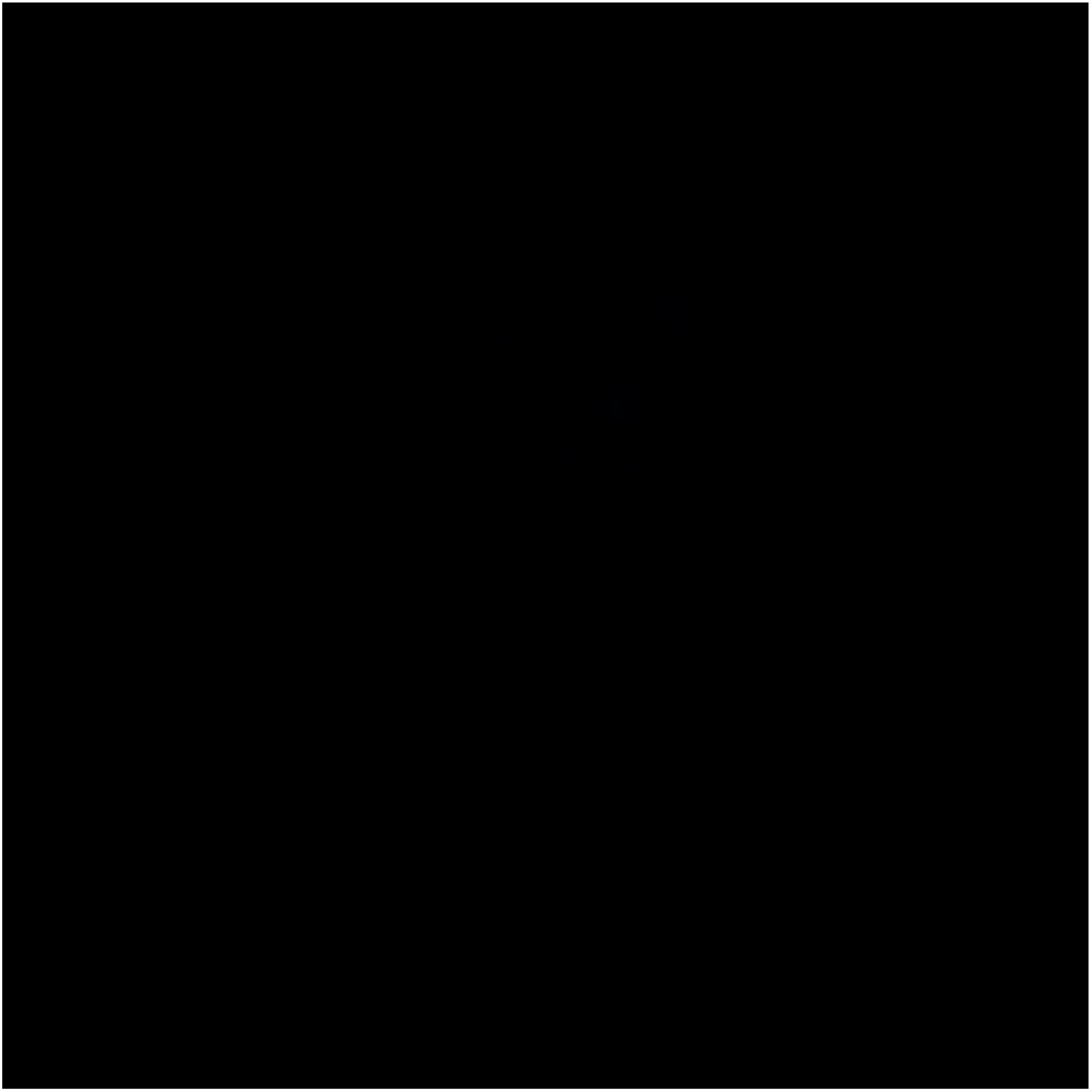
LSM z-stack of a nHDF cell before microchannel

**Supplemental video 2:**
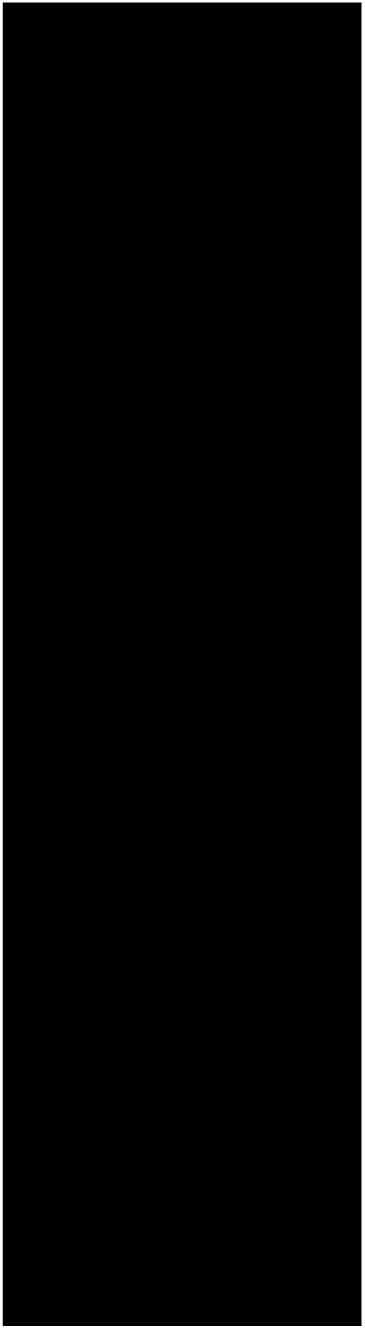
LSM video of a nHDF cell inside a 3 µm microchannel

**Supplemental figure 1:**
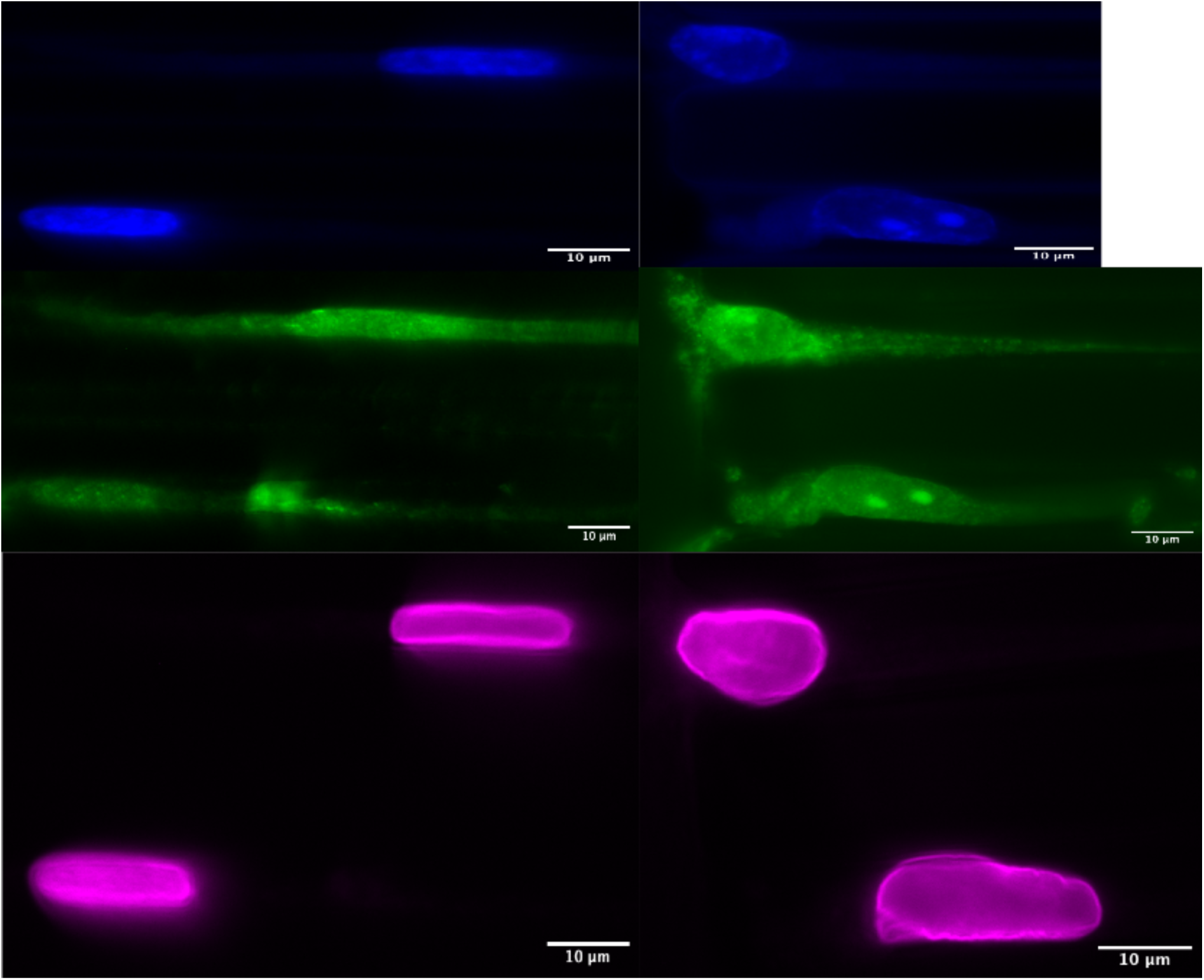
nHDF cells inside microchannels. Left side: nHDF cells inside 3 µm channels. Right side: nHDF cells inside 10 µm channels. Upper row: nHDF cells stained for HOECHST; middle row: CHMP4B staining of nHDF cells. Lower row: lamin A/C staining of nHDF cells

**Supplemental figure 2:**
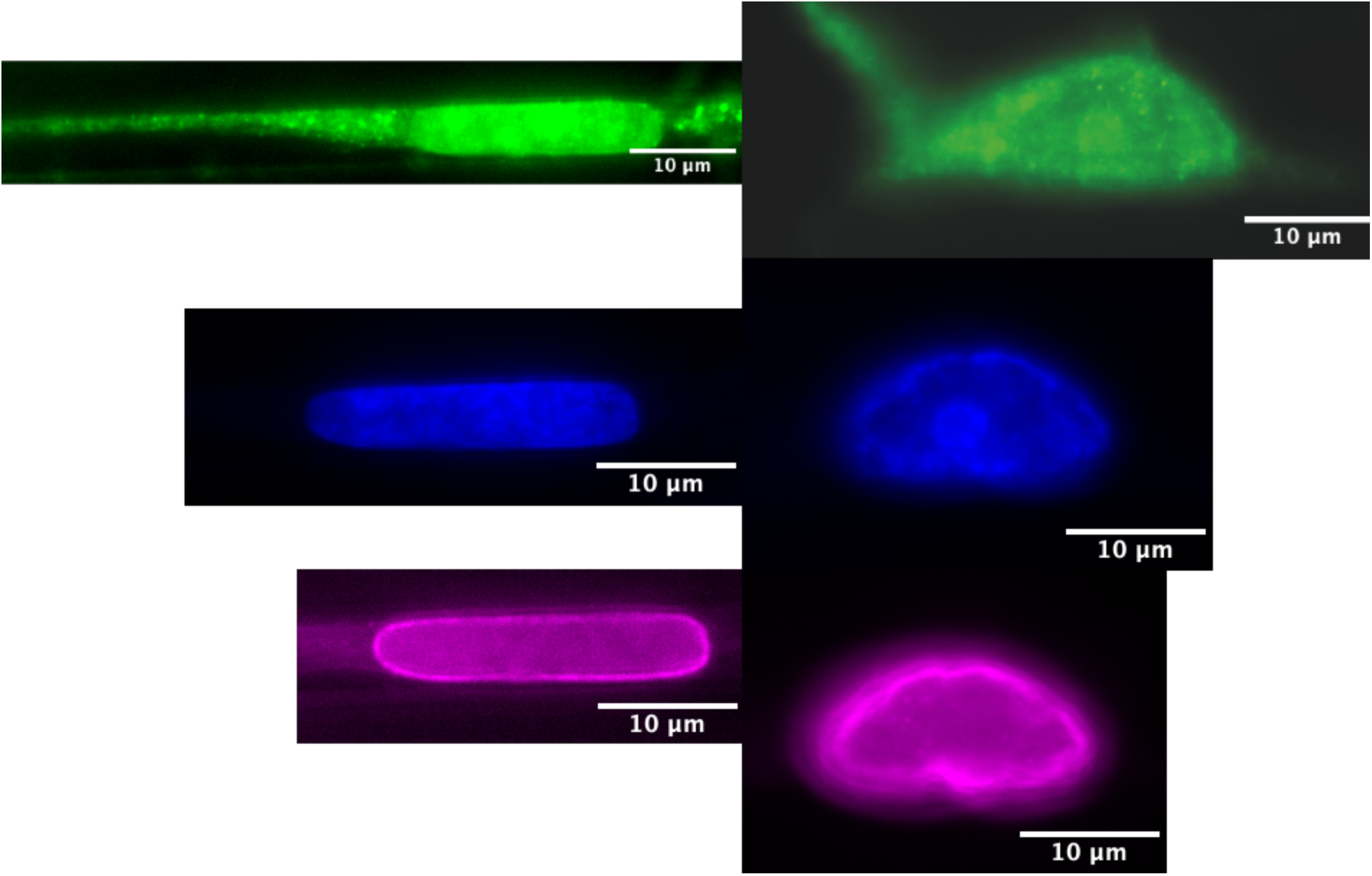
HaCaT cells in microchannels. Left side: HaCaT cells inside 3 µm channels. Right side: HaCaT cells inside 10 µm channels. Upper row: CHMP4B stained HaCaT cells, middle row: HOECHST stained HaCaT cells, lower row:lamin A/C stained HaCaT cells.

**Supplemental figure 3:**
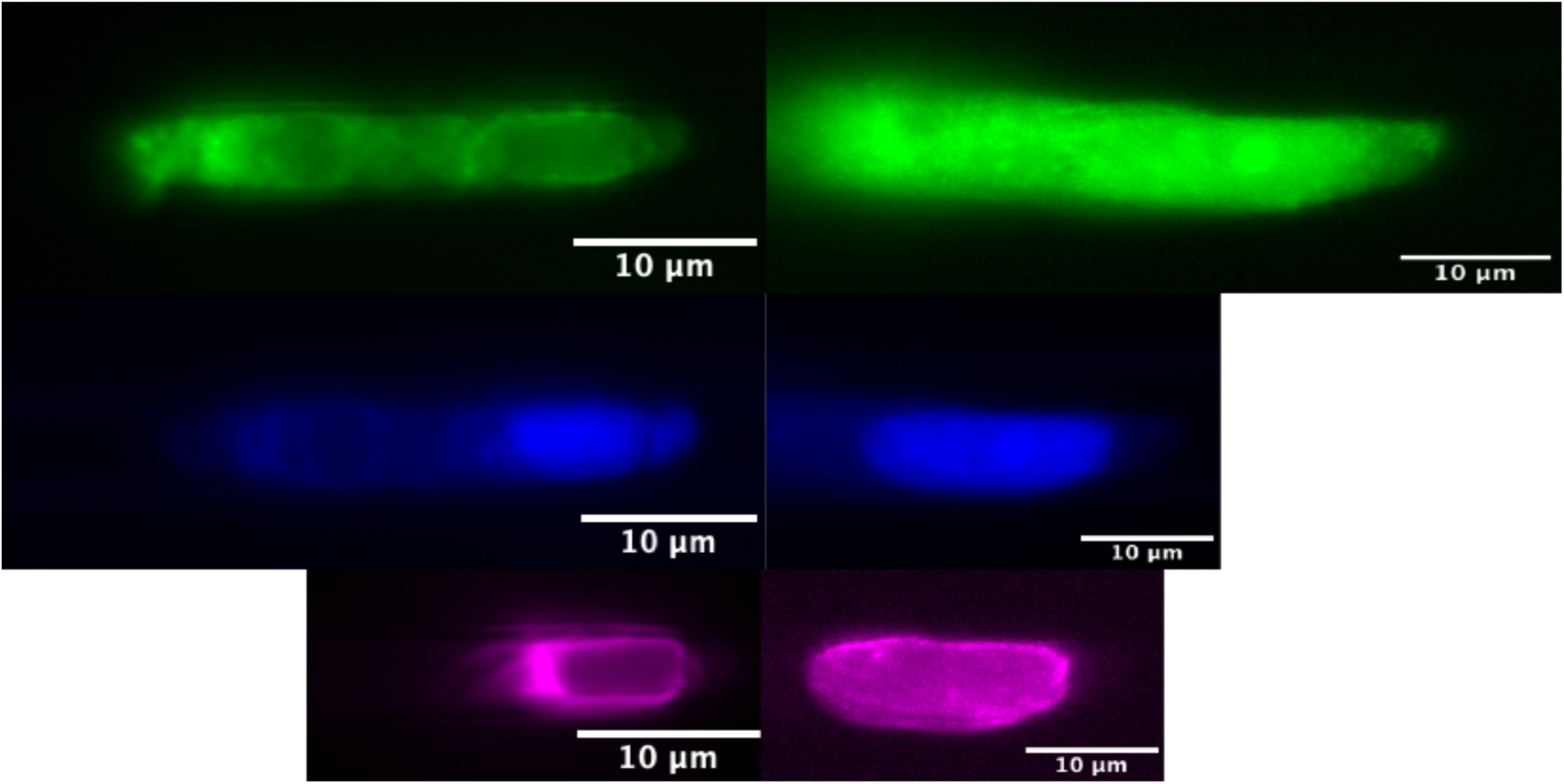
MCF-7 cells in microchannels. Left side: MCF-7 cells inside 3 µm channels, right side: MCF-7 cells inside 10 µm channels. Upper row: CHMP4B stained MCF-7 cells, middle row HOECHST staining of MCF-7 cells, lower row: lamin A/C stained MCF-7 cells.

**Supplemental figure 4:**
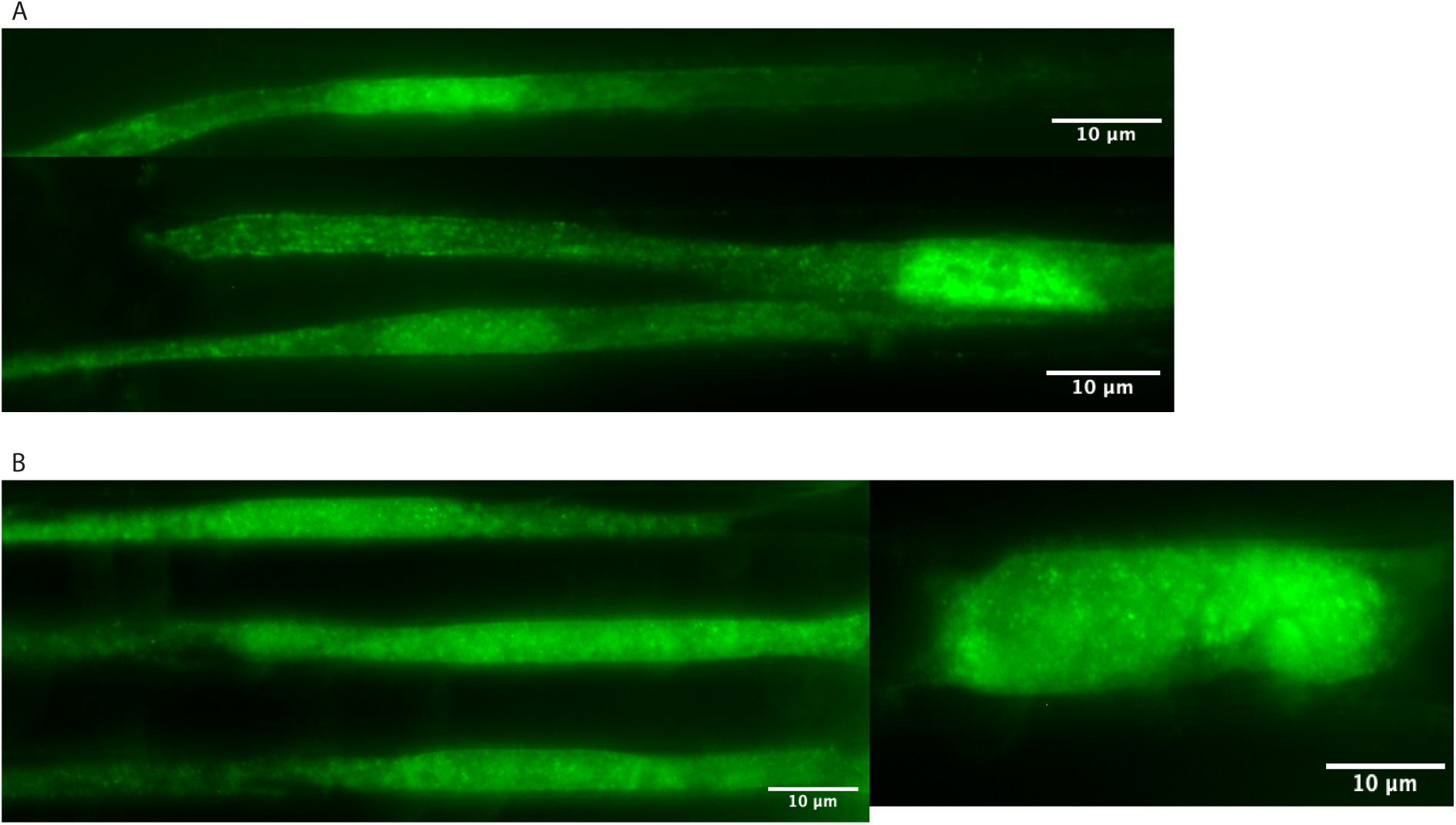
53BP1 staining inside microchannels. A: Upper picture: nHDF cell inside a 3 µm microchannel. Lower picture: nHDF cells inside 10 µm channels. B: Left picture: HaCaT cells inside 3 µm channels, right picture: HaCaT cell inside a 10 µm channel.

**Supplemental video 3:**
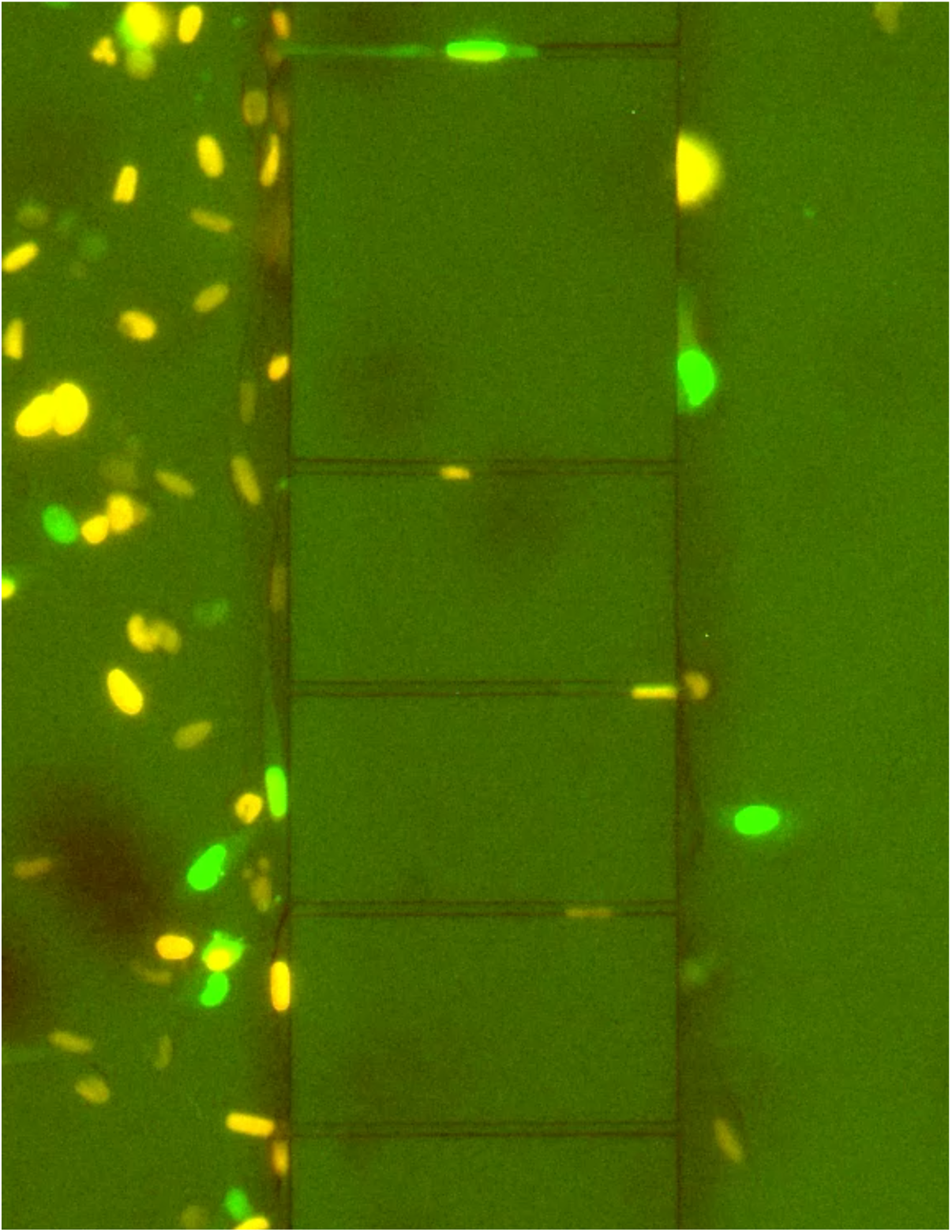
MDA-MB-232 FUCCI cells during their migration through 3 µm channels

**Supplemental video 4:**
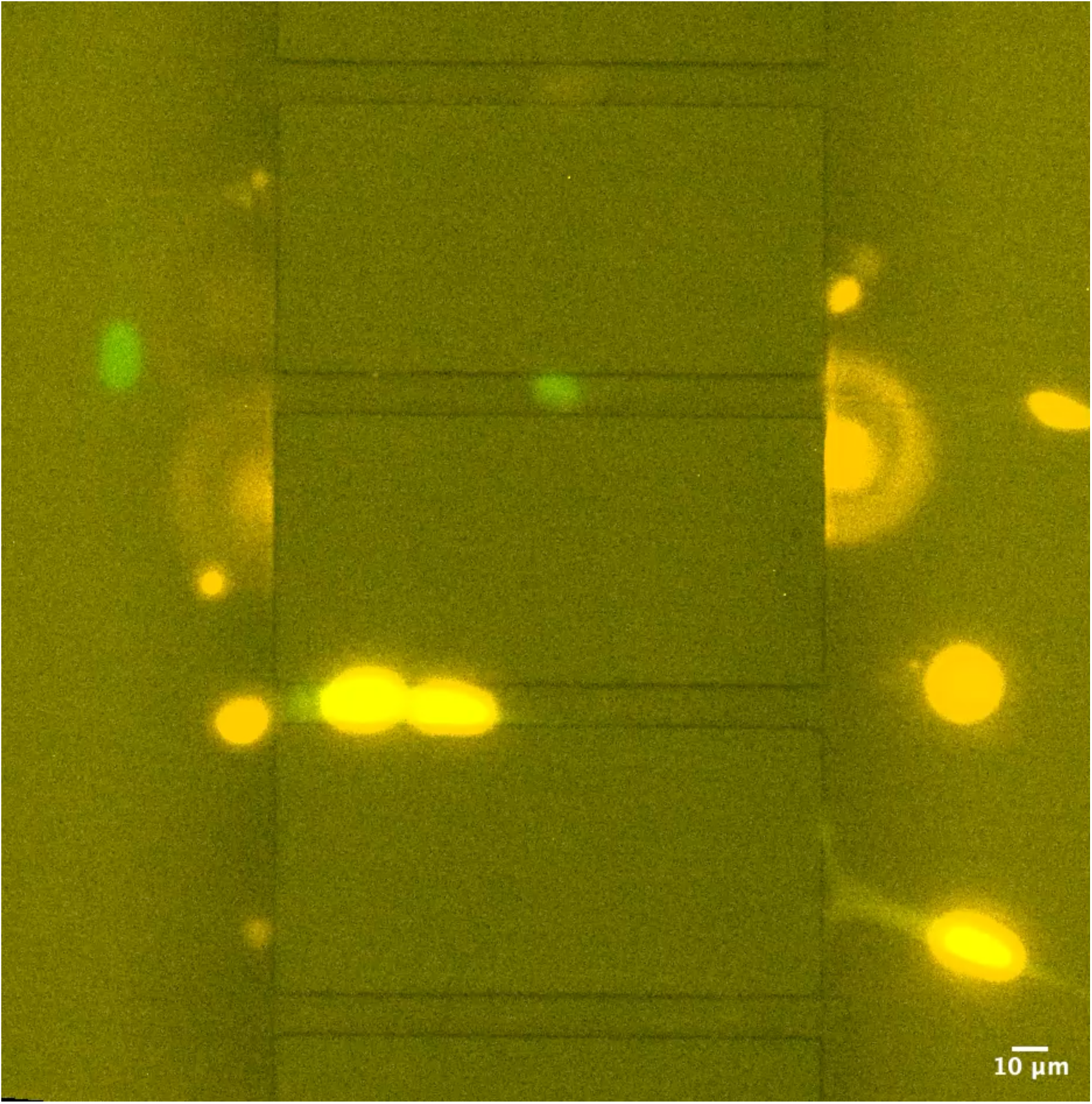
MDA-MB-231 FUCCI during their migration through 10 µm channels

**Supplemental video 5:**
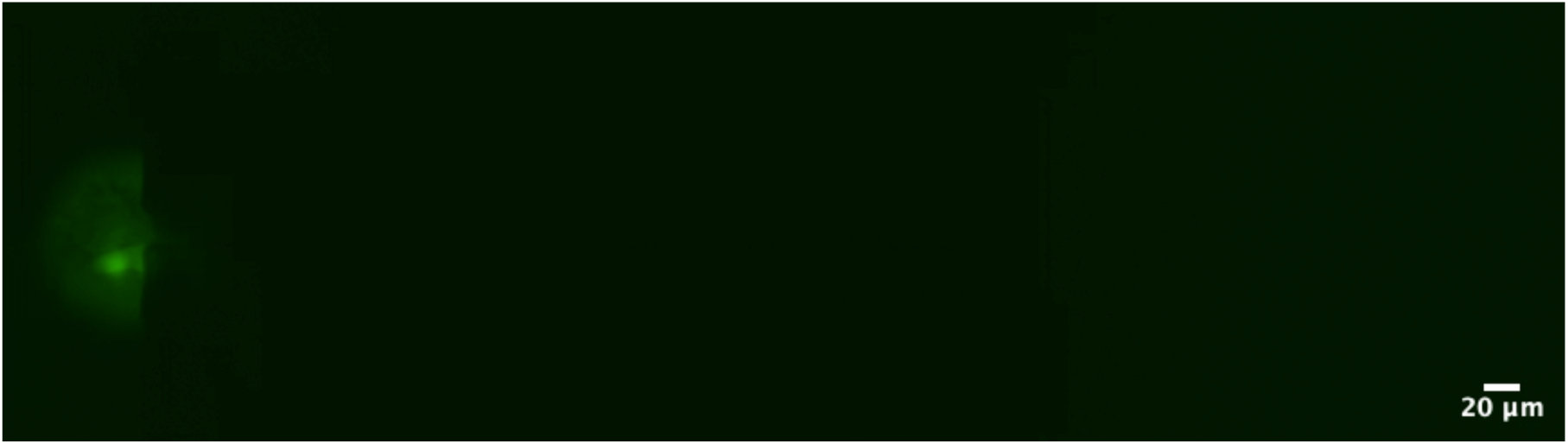
MDA-MB-231 nls-eGFP inside 3 µm channel

**Supplemental video 6:**
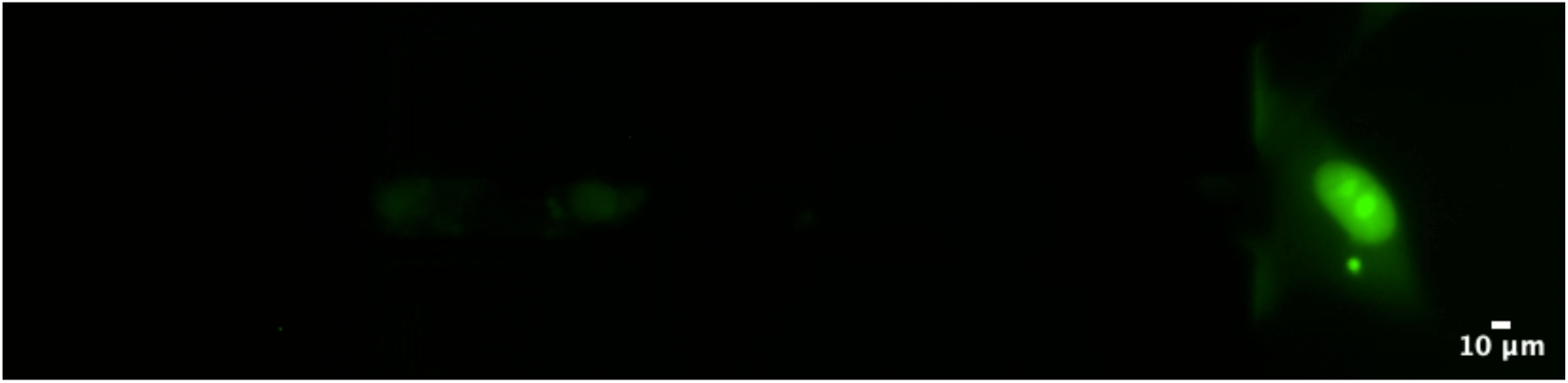
MDA-MB-231 nls-eGFP inside 10 µm channel

