## Supplementary figures and images for "The role of disease state in confined migration"

### Supplemental figure 1

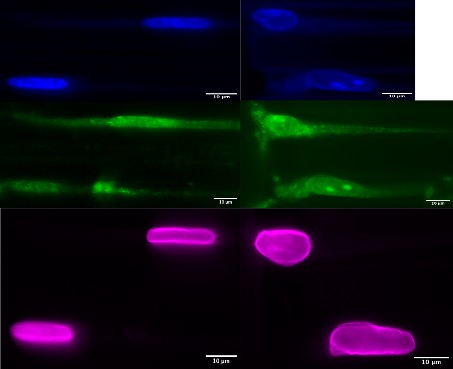

### Supplemental figure 2

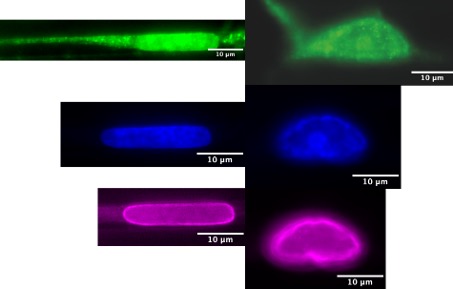

### Supplemental figure 3

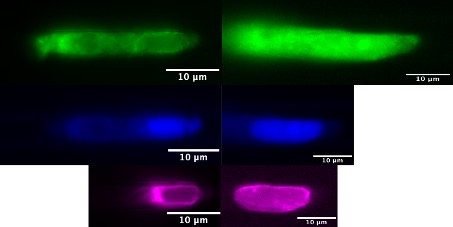

### Supplemental figure 4

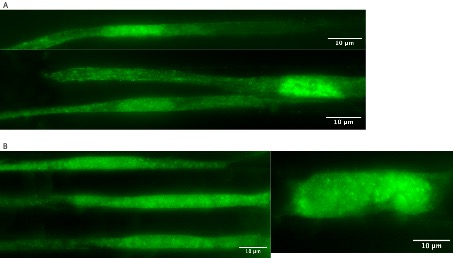
